# Deprivation of dietary fiber enhances susceptibility of mice to cryptosporidiosis

**DOI:** 10.1101/620203

**Authors:** Bruno César Miranda Oliveira, Katia Denise Saraiva Bresciani, Giovanni Widmer

## Abstract

Based on our initial observations showing that mice consuming a probiotic product develop more severe cryptosporidiosis, we investigated the impact of other dietary interventions on the intracellular proliferation of *Cryptosporidium parvum* and *C. tyzzeri* in the mouse. Mice were orally infected with oocysts and parasite multiplication measured by quantifying fecal oocyst output. High-throughput sequencing of 16S ribosomal RNA amplicons was used to correlate oocyst output with diet and with the composition of the intestinal microbiota. On average, mice fed a diet without fiber (cellulose, pectin and inulin) developed more severe infections. As expected, a diet without fibers also significantly altered the fecal microbiota. Consistent with these observations, mice fed a prebiotic product sold for human consumption excreted significantly fewer oocysts. The fecal microbiota of mice consuming no plant polysaccharides was characterized by a lower relative abundance of Bacteroidetes bacteria. Since bacterial metabolites play an important role in the physiology of intestinal enterocytes, we hypothesize based on these observations that the impact of diet on parasite proliferation is mediated primarily by the metabolic activity of the anaerobic microbiota, specifically by the effect of certain metabolites on the host. This model is consistent with the metabolic dependence of intracellular stages of the parasite on the host cell. These observations underscore the potential of dietary interventions to alleviate the impact of cryptosporidiosis, particularly in infants at risk of recurrent enteric infections.

**AUTHOR SUMMARY:** The infection with *Cryptosporidium* parasite, a condition known as cryptosporidiosis, is a common cause of infant diarrhea in developing countries. We have previously shown that mice infected with *C. parvum*, one of the main cause of human cryptosporidiosis, develop a more severe infection if given probiotics. To investigate the mechanism of this effect, we fed mice prebiotics and diet lacking plant fiber. We found that fermentable fiber, whether administered as a prebiotic supplement or is part of the diet, has a protective effect against cryptosporidiosis in mice. We also observed a significant association between the severity of infection and the composition of the gut microbiota. A significant inverse correlation was found between severity of cryptosporidiosis and the ratio between the abundance of bacteria belonging to the phylum Bacteroidetes and the abundance of Firmicute bacteria. This ratio is frequently viewed as a marker of a healthy microbiota. These results raise the possibility that dietary interventions could be used to alleviate the impact of cryptosporidiosis.

## INTRODUCTION

Protozoa of the genus *Cryptosporidium* are important pathogens causing diarrhea in humans, ruminants and other species of animals worldwide [1]. Various *Cryptosporidium* species are recognized as opportunistic pathogens in patients with AIDS, where cryptosporidiosis can lead to protracted diarrhea and wasting. Although immunocompetent patients heal spontaneously within a few weeks, recent studies in developing nations have pointed to *Cryptosporidium* as the second leading cause of infant diarrhea [2, 3].

The resistance of *Cryptosporidium* parasites to anti-protozoal drugs [4], and the lack of alternative therapeutic options, led us to investigate the interaction between the gut microbiota and the parasite. The previously reported unexpected observation that a probiotic product can aggravate the course of cryptosporidiosis in mice [5] supports the hypothesis that parasite proliferation is impacted by diet and possibly by the effect of diet on the gut microbiota. This observation is significant because it could lead to the development of simple dietary supplements for mitigating cryptosporidiosis and perhaps other enteric infections in vulnerable infants.

The benefits to intestinal health of diets rich in plant fibers are well known [6]. It has been suggested that consumption of fiber below nutritional recommendations [7, 8] may lead to dysbiosis. A decrease in the Bacteroidetes/Firmicutes ratio has often been linked to a poor intestinal health index and to obesity [9]. Dysbiosis may also deplete the intestinal mucosal layer [10]. To what extent mucus depletion may play a role in susceptibility to cryptosporidiosis has not been investigated. Several mechanisms linking diet, microbiota and enteric infections have been proposed [11]. Bacterial metabolites, particularly those originating from the fermentation of certain plant polysaccharides, have been shown to play and important role in modulating the resistance to enteric bacterial infections [12]. Research on the interaction between the microbiota and the intestinal epithelium has shown the importance of bacterial metabolites, such as short-chain fatty acids originating from the anaerobic breakdown of plant polysaccharides [10]. The role of the intestinal microbiota in regulating the immune response and preventing inflammation has also been investigated [11, 13]. With respect to enteric infections, much research has focused the protective role of the microbiota, a phenomenon often referred to as “colonization resistance” [14, 15]. In contrast to what is known about the effect of diet and bacterial metabolites on the intestinal physiology, less research has focused on mechanisms linking diet and enteric infections. This limitation is particularly true for enteric protozoa [16].

With respect to cryptosporidiosis, research with germ-free severe combined immunodeficient (SCID) mice and SCID mice colonized with intestinal microbes conducted by Harp and co-workers showed that a normal intestinal microbiota delayed the onset of *C. parvum* oocyst excretion by several weeks [17, 18]. A protective role of the gut microbiota against cryptosporidiosis was also observed in neonatal mice [19, 20] A protein-deficient diet was also found to increase susceptibility of mice to *C. parvum* [21]. This phenotype was attributed to a reduced epithelial cell turnover. The effect of probiotics on the course of cryptosporidiosis was also observed by others [22, 23]. This research uncovered a beneficial effect of *Enterococcus faecalis* administration to mice infected with *C. parvum*. None of these studies have investigated potential mechanisms mediating the observed effect on the development of *C. parvum*.

Here we describe experiments with a mouse model of cryptosporidiosis aimed at investigating changes in the bacterial microbiome caused by dietary fiber and at relating these changes to the severity of cryptosporidiosis. The results show that relatively small changes in diet, or the administration of a prebiotic formulation, can reduce the severity of cryptosporidiosis.

## MATERIAL AND METHODS

### Parasites

*C. parvum* strain TU114 oocysts [24] was used in experiment 1 and 4 whereas *C. tyzzeri* oocysts were used in experiments 2, 3 and 5. *C. parvum* strain TU114 belongs to the anthroponotic subgroup characterized by a GP60 surface glycoprotein genotype IIc [25, 26]. *C. tyzzeri* is a species commonly found in domestic mice of the species *Mus musculus* [27]. Oocysts for the experimental infections were purified from feces of mice on Nycodenz (Alere Technologies, Oslo, Norway) step gradients as previously described [28]. The age of the oocysts was 65, 37, 22, 38 and 13 days for experiments 1, 2, 3, 4 and 5, respectively (Table 1).

**Table 1.** Summary of experiments.

| Experiment | Isolate | Groups | Mouse Strain | Treatment | Dex<sup>1</sup><br>treatment | Oocyst age<br>(days) | Number of<br>mice |
| --- | --- | --- | --- | --- | --- | --- | --- |
| 1 | TU114 | 2 | CD-1 | no-fiber diet | 16 mg/l | 65 | 8 |
| 2 | *C. tyzzeri* | 2 | C57BL/6 | no-fiber diet | - | 37 | 8 |
| 3 | *C. tyzzeri* | 4 | C57BL/6 | no-fiber diet | - | 22 | 12 |
| 4 | TU114 | 4 | CD-1 | prebiotics | 16 mg/l | 38 | 16 |
| 5 | *C. tyzzeri* | 4 | C57BL/6 | prebiotics | - | 13 | 12 |

<sup>1</sup> Dexamethasone concentration in drinking water

### Mouse experiments

To test the effect of dietary fiber, three experiments were performed using no-fiber diet and matched control diet (“medium-fiber diet”) (Supplementary Table 1). In experiment 1, 8 female CD-1 mice aged ∼5 weeks were randomly divided into two groups and immunosuppressed by adding disodium dexamethasone 21-phosphate (Sigma, cat. D1169) to drinking water at a concentration of 16 mg/L [29]. The immunosuppressive treatment was initiated on the day −5 post-infection (PI), where day 0 is the day of infection. In experiment 2, we used 8 female C57BL/6 mice, also divided into two groups of 4 mice. In experiment 3, 12 female C57BL/6 mice were divided into four groups of 3 mice. In all experiments, mice were provided *ad libidum* with autoclaved water. In experiments 1 and 2, each group was fed one type of diet and in experiment number 3, two groups ingested medium-fiber diet and two groups no-fiber diet. The diet was given starting on day −5 PI, i.e., 5 days before the animals were infected with *Cryptosporidium* oocysts. To test the effect of prebiotics on the microbiome and on the excretion of *Cryptosporidium* oocysts, we performed two experiments. In experiment 4, 16 CD-1 mice, randomly divided into 4 groups of 4 mice, were given normal diet and were immunosuppressed by the addition of dexamethasone to drinking water at a concentration of 16 mg/L. In addition to immunosuppression with dexamethasone, vancomycin and streptomycin were added to drinking water at a concentrations of 500 mg/L and 5 g/L, respectively, starting on day −6 PI. Metronidazole at the dose of 20 mg/kg was given daily by gavage, starting at day 6 PI. Antibiotic treatment was terminated on day 2 PI. The goal of the antibiotic treatment was to deplete the native intestinal microbiome [30], and replicate the treatment used in a previous series of experiments with probiotics [5]. From day −1 PI, the drinking water was supplemented with prebiotic (Supplementary Table 1) at a concentration of 2.8 g/L. Lastly, in experiment 5, 12 immunocompetent C57BL/6 mice divided into four groups were used, two were given prebiotic in the drinking water starting on day −5 PI, and the other two groups drank unsupplemented water. In this experiment all groups ingested medium-fiber diet. Experiments typically lasted 3 weeks.

Upon arrival, each mouse was individually tagged and randomly assigned to a treatment groups (Table 1). Mice were orally infected on day 0 PI with approximately 2 × 10^4^ oocysts of *C. parvum* strain TU114 (experiment 1 and experiment 4) or *C. tyzzeri* (experiment 2, 3 and 5). To obtain fecal pellets for intestinal microbiota analysis, mice were individually transferred to a 1-L plastic cup and fecal pellets collected immediately after defecation. The pellets were stored at −20°C. To collect feces for oocyst enumeration using flow cytometry, mice were individually transferred overnight to collection cages fitted with a wire bottom. Feces collected overnight were stored at 4°C. Following overnight fecal collection, mice were returned to regular cages with their original cage mates. On days when feces were collected for oocyst enumeration, mice were individually housed for 14-16 h and spent the remaining time in regular cages with their respective cage mates.

### Oocyst enumeration

Prior to processing for flow cytometry (FCM), fecal pellets were suspended in water and homogenizing to a slurry. The water volume was adjusted according the the volume of feces and varied between 1.5 ml and 4 ml. A previously described procedure [5] was used to fluorescently label oocysts. The only modification consisted in filtering the fecal slurries through a 38-μm opening Nylon mesh, (Component Supply, Sparta, Tennessee, cat. 06725-01) before FCM. For each experiment, three samples were randomly selected for replication. Replication involved the processing and labeling of 5 separate aliquots originating from a fecal sample. The labeled samples were analyzed by FCM using a Becton Dickinson Accuri C6 cytometer. Oocyst counts were normalized against the volume of fecal slurry.

### Microbiota analysis

The procedures for DNA extraction, amplicon library construction and bioinformatics were previously described [5, 31]. Briefly, fecal DNA was PCR amplified to prepare amplicons of the V1V2 variable region of the bacterial 16S rRNA gene [32, 33]. The multiplexed amplicon library was size-selected on a Pippin HT system (Sage BioScience, Beverly, Massachusetts) and sequenced in an Illumina MiSeq sequencer at the Tufts University genomics core facility (tucf.org) using single-end 300 nucleotide strategy. To control for technical variation introduced during PCR, library preparation and sequencing, each library included two replicates of two randomly selected samples. Replication involved the separate processing of duplicated fecal samples and tagging each amplicon with a different barcode.

### Bioinformatics

FASTQ formatted sequences were processed using programs found in *mothur* [34] essentially as described [5, 35]. Briefly, random subsamples of 5000 sequences per sample were processed. Pairwise UniFrac phylogenetic distances [36] between samples were calculated in *mothur*. Analysis of Similarity (ANOSIM) [37] was used to test the significance of clustering by treatment. Program *anosim* was run in *mothur* using a weighted UniFrac distance matrix as input. Operational Taxonomic Units (OTUs) were obtained using program *cluster*, using the OptiClust clustering method [38]. A distance cut-off of 3% was applied.

Redundancy Analysis (RDA) was used to test the significance of association between OTU profile and oocyst concentration. The program was run in CANOCO [39]. The pseudo-F statistic was calculated by Monte Carlo with 1000 permutations of samples between treatment groups. OTU abundance values for the 150 most abundant OTUs served as dependent variables. Oocyst concentration determined by flow cytometry as described above served as independent variable. Where two experiments were pooled, i.e., experiments 2 and 3, any effect of the experiment was excluded by defining the experiment as covariate. GenAlEx [40] was used to draw Principal Coordinate Analysis (PCoA) plots using weighted UniFrac distance matrices as input. Linear Discriminant Analysis as implemented in program LEfSe [41] was used to identify statistically significant differences in OTU abundance profiles between two groups of samples defined by the dietary treatment.

### Sequence data accession numbers

Sequence data from experiments 1-5 were deposited in the European Nucleotide Archive under study accession numbers PRJEB31954, PRJEB31955, PRJEB31958, PRJEB31959 and PRJEB31960, respectively.

### Ethics statement

The animal experiments adhered to the National Institutes of Health’s Public Health Service Policy on Humane Care and Use of Laboratory Animals. The animal experiments were approved by the Tufts University Institutional Animal Care and Usage Committee (IACUC). The IACUC approved the experiments described above and as described in document G2016-40.

## RESULTS

### Effect of diet and dietary supplements on the course of cryptosporidiosis

To test whether no-fiber diet affects the severity of cryptosporidiosis, immunosuppressed and competent mice were infected with *C. parvum* and *C. tyzzeri* oocysts, respectively. The intensity of the infection in mice fed medium-fiber diet or regular diet was measured by quantifying oocyst output by FCM 6 times over the duration of each experiment. In the 3 experiments designed to compare the effect of dietary fiber, a significant increase in oocyst excretion by mice ingesting no-fiber diet was detected (Table 2). Fig. 1 shows overnight oocyst output over time for the three experiments. We performed analogous experiments to test the effect of prebiotics, which are in essence fermentable fibers (Supplementary Table S1). In experiment 4, prebiotics were given to mice fed medium-fiber diet. The impact on the severity of *Cryptosporidium* infection was also significant (Table 2). Similarly, giving prebiotics to mice fed no-fiber diet (experiment 5) significantly reduced oocyst output over the course of the infection (Table 2). Fig. 2 shows the pattern of oocyst production over time for prebiotic experiments 4 and 5.

**Table 2.** Results of oocyst flow cytometry enumeration and 16S amplicon sequence analysis.

| Experimenttreatment | Mann-Whitney (FCM) |  |  |  |  | ANOSIM (microbiota all days) |  |  |  |
| --- | --- | --- | --- | --- | --- | --- | --- | --- | --- |
|  | Number<br>of fecal<br>samples | Mean<br>treatment<sup>2</sup> | Mean<br>control | *U* | *P* | Number<br>of fecal<br>samples | Comparison | *R* | *P* |
| 1 | 30 | 8058 | 1023 | 56 | 0.020 | 31 | nfd - mfd<sup>1</sup> | 0.287 | < 0.001 |
| 2 | 48 | 3069 | 1265 | 118 | 0.001 | 32 | nfd - mfd | 0.710 | < 0.001 |
| 3 | 72 | 2634 | 1536 | 392 | 0.004 | 50 | nfd - mfd | 0.495 | < 0.001 |
| 4 | 95 | 552 | 3201 | 542 | <<br>0.001 | 50 | Prebiotic, mfd | 0.059 | 0.046 |
| 5 | 60 | 566 | 936 | 305 | 0.033 | 49 | Prebiotic, nfd | 0.202 | < 0.001 |

<sup>1</sup>nfd, no-fiber diet; mfd, medium-fiber diet
<sup>2</sup>Treatments, see Table 1

**Fig. 1.**
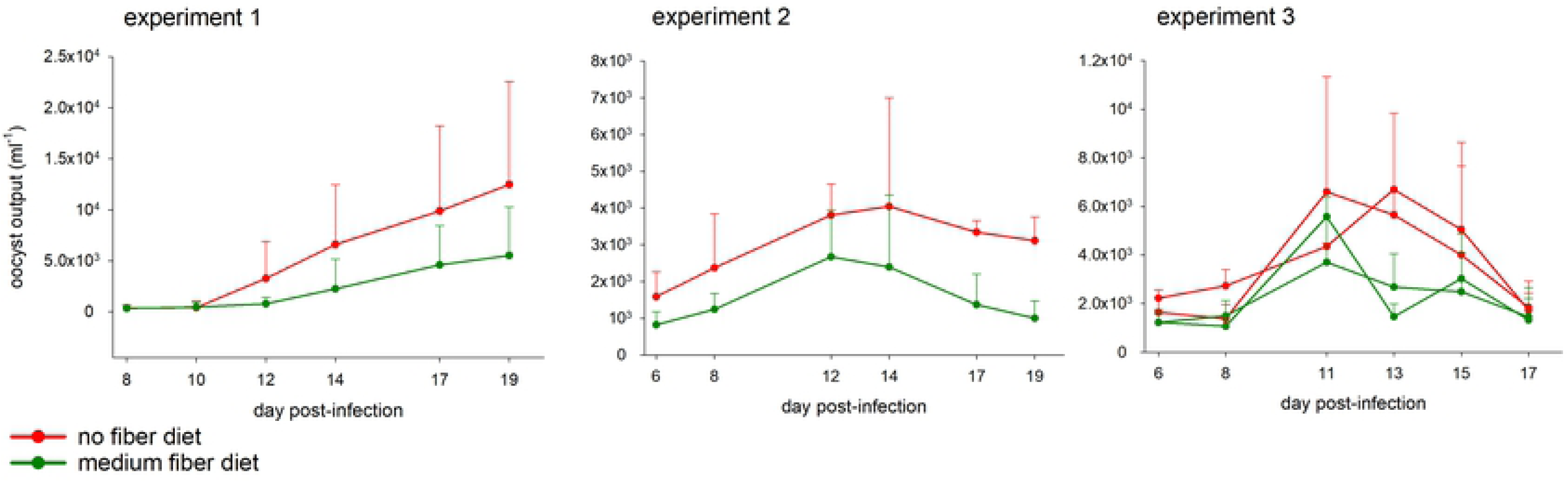
Effect of diet on severity of *C. parvum* and *C. tyzzeri* infection. The Y axes show mean overnight oocyst output per ml fecal slurry by group. The mice were sampled individually over a period of approximately 16 h. Values represent mean of 3 mice per group. Experiment 1, *C. parvum*; Experiment 2, *C. tyzzeri*; Experiment 3, *C. tyzzeri*. See Table 1 for additional details. Error bars show standard deviation (SD). Color indicates treatment as indicated. One mouse from experiment 1, no-fiber group, remained uninfected for the duration of the experiment and was excluded.

**Fig. 2.**
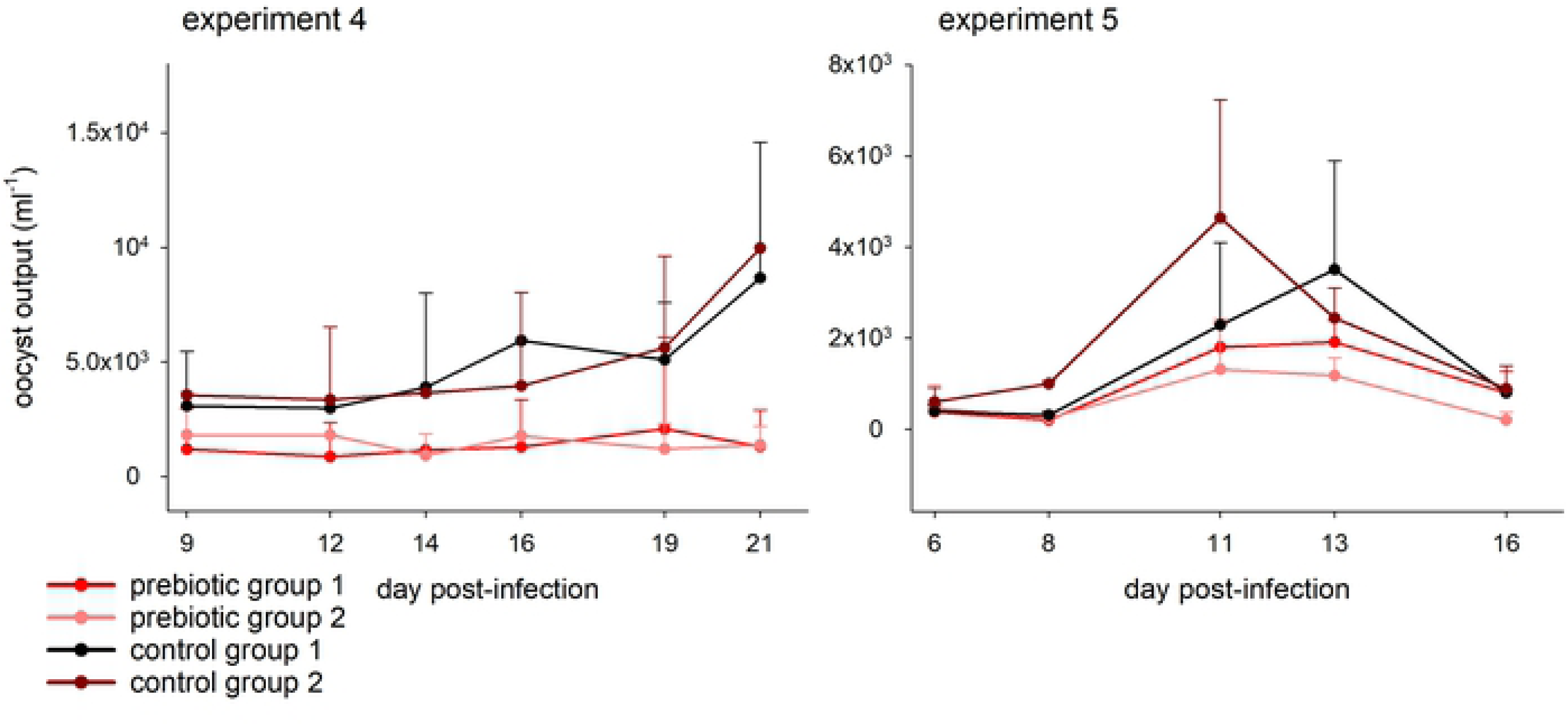
Effect of prebiotic on severity of *C. parvum* and *C. tyzzeri* infection. Datapoints represent mean of 4 mice (experiment 4) and 3 mice (experiment 5). Error bars show SD. Color indicates treatment as follows: red and pink, supplemented with prebiotic; black and brown, control. See Fig. 1 legend, Table 1 and Materials and Methods for details.

Based on the results described above and on previously published observations [5], we investigated whether the effect of diet and dietary supplements on cryptosporidiosis could be mediated by the intestinal microbiota. To evaluate this model, the fecal bacterial microbiota was analyzed using 16S amplicon sequencing. Weighted UniFrac distances [36] between pairs of microbiota from each experiment were visualized on PCoA plots (Fig. 3, 4). In experiments 1, 2 and 3, (no-fiber vs. medium-fiber diet), fecal sample collection was initiated on the fifth day after the onset of dietary intake, the day the mice were infected, and continued until day 23 of treatment (day 18 PI). Feces were collected four times during this interval. Demonstrating an effect of diet on the intestinal microbiota, this analysis revealed a non-overlapping distribution of data points according to dietary treatment. ANOSIM *R*-values between diet groups for the three experiments testing the effect of diet are statistically significant (Table 2). Significant clustering according to prebiotic treatment was observed in experiments 4 and 5 based on 50 and 49 samples, respectively, collected between day 0 PI and day 15 PI. Consistent with a significant effect of the prebiotics, the ANOSIM *R*-value in experiment 4 was 0.059, (p = 0.046) and in experiment 5 0.202 (p < 0.0001). (Table 2).

**Fig. 3.**
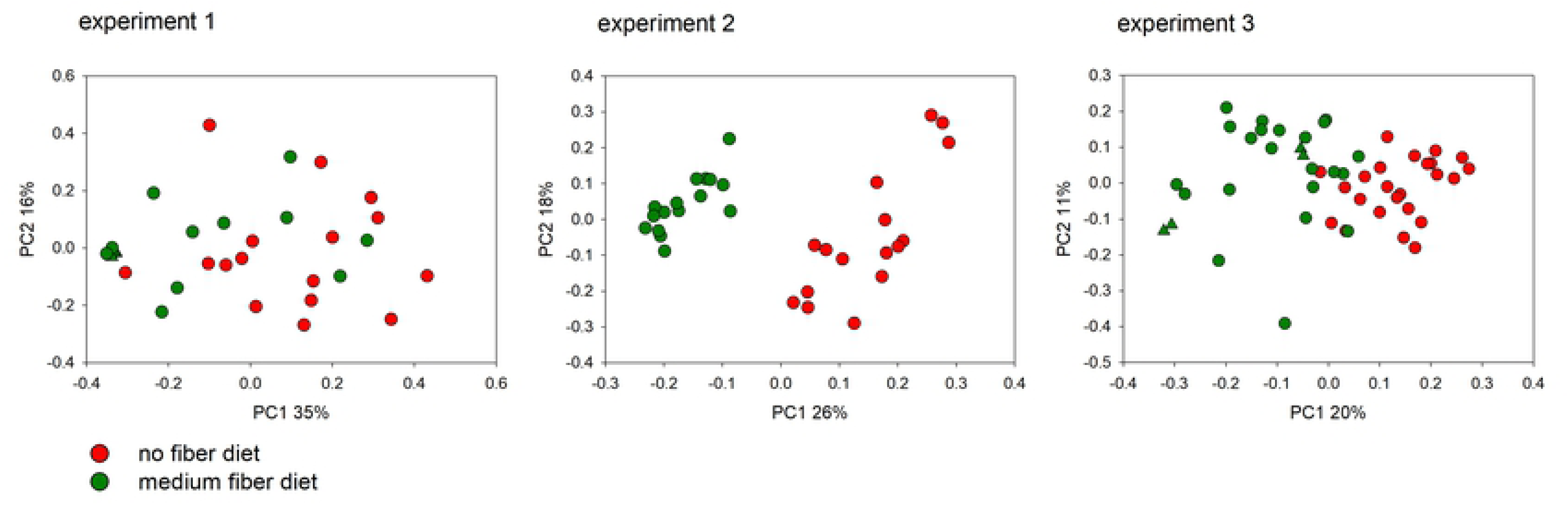
Impact of diet on the fecal microbiome of mice infected with *C. parvum* (experiment 1) and *C. tyzzeri* (experiment 2/3). Principal Coordinate Analysis was used to display weighted UniFrac distances between pairs of fecal microbiota. Experiment 1 analysis includes data from 31 fecal samples collected from individual mice starting on day 5 of treatment (day 0 PI) until day 21 of treatment (day 16 PI). For experiment 2/3, 32 and 50 samples, respectively, from individual mice were analyzed. Each data point represents one fecal sample, color-coded according to treatment. Matching triangle symbols indicate replicate analyses of the same fecal samples.

**Fig. 4.**
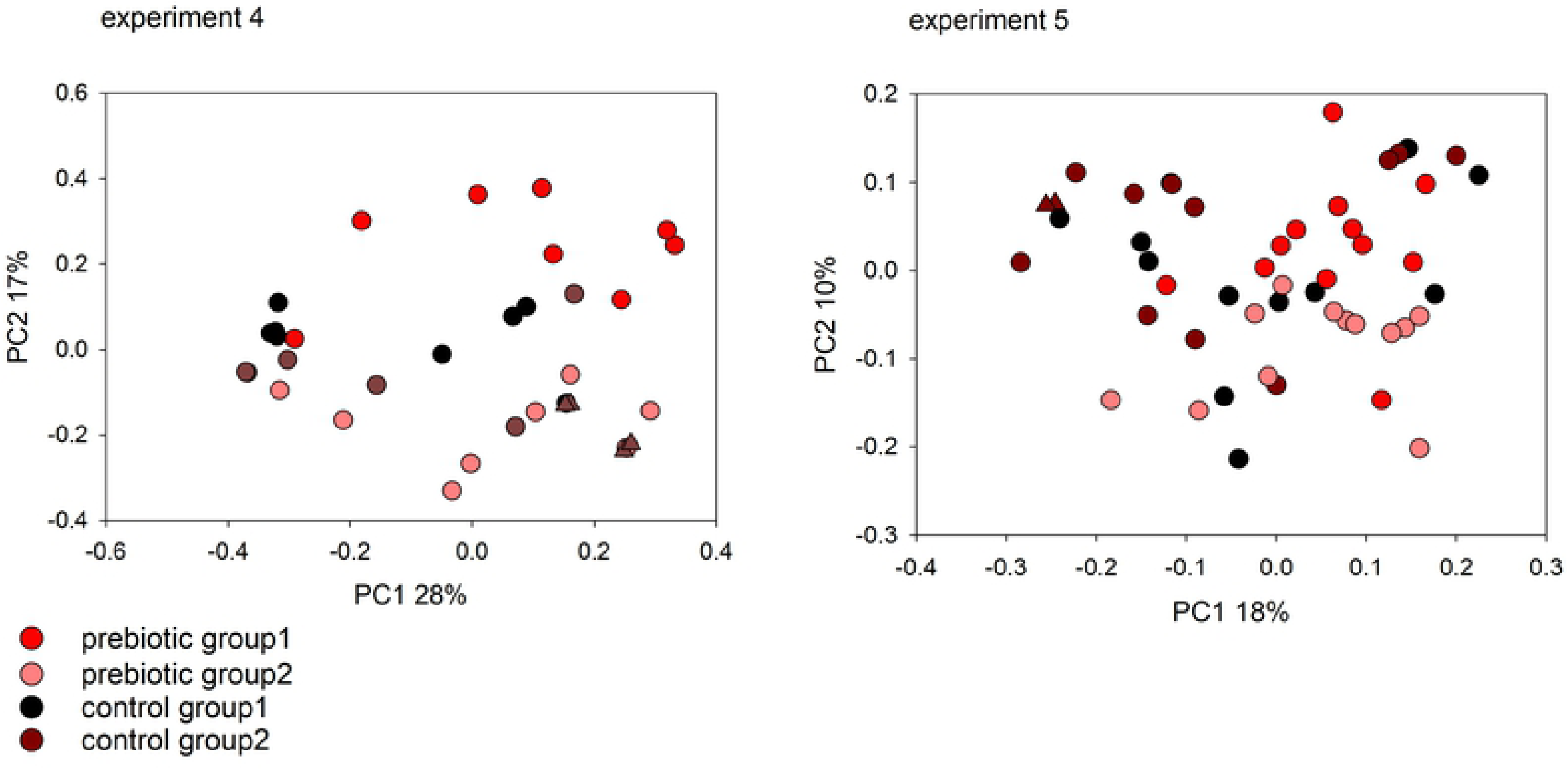
Impact of diet on the fecal microbiome of *C. parvum* (experiment 4) and *C. tyzzeri* (experiment 5) infected mice. Principal Coordinate Analysis was used to display weighted UniFrac distances between pairs of fecal microbiome samples. Experiment 4 analysis includes data from 50 fecal samples. For experiment 5, 49 samples were analyzed from individual mice. Each data point represents one sample, color-coded according to treatment and group as shown in Fig. 2. Matching triangle symbols indicate replicate analyses of the same fecal samples.

### Diet and oocyst output

Having detected an association between dietary fiber and cumulative oocyst output, and dietary fiber and fecal microbiota profile, we focused on the microbiome on day 0 PI. We reasoned that if the gut microbiota impacts the severity of cryptosporidiosis, the microbiota on day 0 PI would be the most relevant to examine. Since colonization of the gut epithelium with *Cryptosporidium* is known to impact the microbiota [35], the analysis of the microbiota on day 0 PI eliminates the effect of cryptosporidiosis on the microbiota and enables detecting any effect of the microbiota on cryptosporidiosis. The effect of no-fiber diet on the fecal microbiota was already detectable after 5 days of treatment (day 0 PI) in experiments 2 and 3 (ANOSIM *R* = 0.82, p = 0.03; *R* = 0.74, p = 0.001, respectively). In experiment 1, the effect was not significant (ANOSIM *R* = 0.17, p = 0.22). Administration of prebiotics in experiments 4 and 5 did not significantly change the microbiota composition according to ANOSIM (*R* = 0.10, p = 0.06, n = 16; *R* = 0.06, p = 0.29, n = 12, respectively). To examine to what extent the day 0 microbiota composition correlates with total oocyst output over the course of the infection, we merged experiments 2 and 3, which are exact replicates, to increase the power of the analysis. Experiment 1 was excluded from this analysis because the mice were immunosuppressed, because the microbiota on day 0 did not show any impact of diet and because of mortality only 31 samples were available. For the remaining 4 experiments, we analyzed the correlation between cumulative oocyst output for each mouse and the microbiota OTU profile using RDA. Of the 20 day 0 microbiota samples from pooled experiments 2 and 3, 8 originated from experiment 2 and 12 from experiment 3. Defining the experiment as covariate, a Monte Carlo permutation test indicated a significant correlation between cumulative oocysts output and the OTU profile (pseudo-F = 2.1, p = 0.0354). As expected from the lack of prebiotic effect on the microbiota on day 0, RDA of experiment 4 and 5 showed a non-significant association between day 0 microbiota profile and cumulative oocyst output (pseudo-F = 0.7, p = 0.4688; pseudo-F = 1.3, p = 0.172, respectively).

### Analysis of bacterial taxonomy

As expected from the different treatments used in the 5 mouse experiments, the taxonomy of the fecal bacterial microbiota differed extensively between experiments. Supplementary Fig. 1A illustrates the magnitude of the effect of diet and antibiotics pretreatment on the microbiota. As expected, pre-treating mice with antibiotics in experiment 4 profoundly modified the microbiota when compared with microbiota from untreated mice. Removing the data points from experiment 4 from the PCoA reveals the impact of dexamethasone treatment and/or *Cryptosporidium* species on the microbiota (Supplementary Fig. 1B). Since the experiments were not designed to investigate the effect of these variables, we cannot infer the relative effect of each of these 2 variables on the microbiota. This is because all *C. parvum* infected mice were immunosuppressed, whereas infection with *C. tyzzeri* does not require immunosuppression. The position of experiment 1 data points in Supplementary Fig. 1B also indicates that immunosuppression and/or parasite species has large effect on the microbiota as compared to diet.

The combined samples collected on day 0 PI from experiments 2 and 3 were the primary focus of a taxonomic analysis because of the relatively large sample size (n = 20 mice). Combining these two experiments is consistent with them being exact replicates (Table 1). LDA, as implemented in program LefSe, was used to identify bacterial taxa significantly associated with dietary treatment. This analysis identified 95 taxa significantly more abundant in the no-fiber microbiota and 92 in the medium-fiber microbiota (Supplementary Table S2). Of the 95 taxa in the former group, only 24 (25%) belonged in the phylum Bacteroidetes, which compares to 42 (45%) Bacteroidetes taxa in the medium-fiber group. A Chi-square test confirms that Bacteroidetes taxa were significantly enriched in mice consuming medium-fiber diet (χ^2^ = 8.5, p = 0.003). Given the wide interest in the Bacteroidetes/Firmicutes ratio as a marker of a healthy gut microbiota [42-44], we calculated day 0 Bacteroidetes/Firmicutes from experiment 2/3. As shown in Fig. 5, cumulative oocyst output was negatively correlated with Bacteroidetes/Firmicutes (Pearson r = −0.47, p = 0.04; Spearman r_s_ = −0.46, p = 0.04). As expected from the metabolic function of the Bacteroidetes microbiota, mean Bacteroidetes/Firmicutes on day 0 PI was also significantly correlated with diet (mean no-fiber diet = 2.213, mean medium-fiber diet = 3.950; Mann-Whitney *U* = 305, p = 0.001). In the other experiments this correlation was not observed on day 0 PI, but calculating the Bacteroidetes/Firmicutes ratio for the entire experiment (all time points), revealed a significant effect of diet in experiment 1 (n = 31, *U* = 160, p = 0.025) and in experiment 5 (n = 49, *U* = 181, p = 0.018) As indicated above, this outcome could however be related to the effect of the infection of the microbiota. In experiment 4, we did not observe a significant difference between treatment groups (n = 50, *U* = 180, p = 0.9). This observation is consistent with the fact that in this experiment a prebiotic supplement given to mice fed medium-fiber diet. In addition, pretreatment of mice with antibiotics in this experiment profoundly impacted the microbiota (Supplementary Fig. 1).

**Fig. 5.**
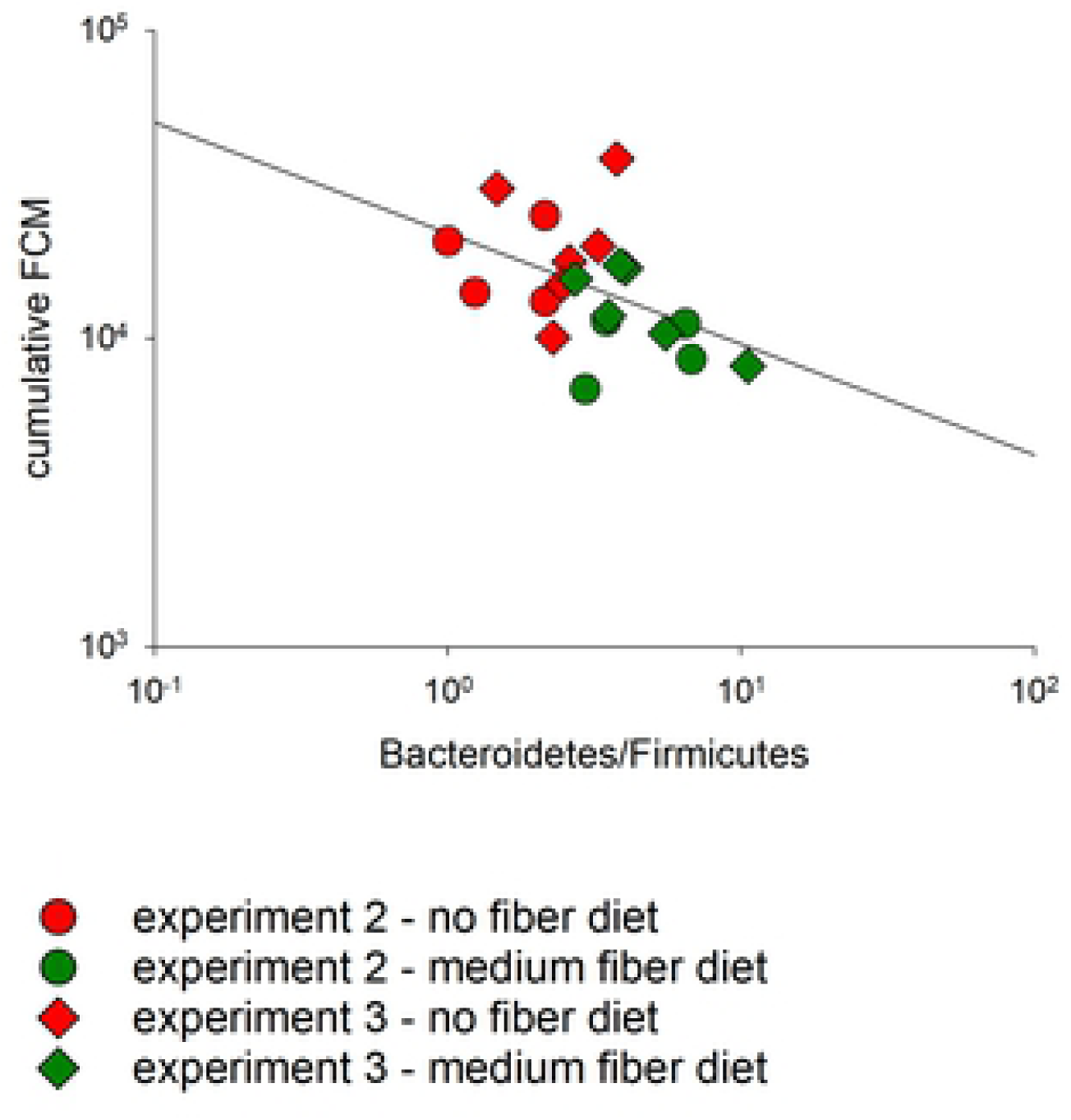
Bacteroidetes/Firmicutes ratio on the day of infection is negatively correlated with total oocyst output. The Bacteroidetes to Firmicutes relative abundance ratio was calculated for 20 samples from day 0 PI of experiments 2 and 3 and plotted against cumulative oocyst output on a log-log plot. 16S sequences de-noised as described in Material and Methods were classified using the method described by Wang et al. [58]. The minimum bootstrap value for taxonomic assignment was set at 70%. Color indicates diet and symbol experiment, as shown in the key.

## DISCUSSION

Consistent with previously published observations [5], the results presented here show that in the mouse a diet low in fermentable fiber content impacts the intestinal microbiota and aggravates the infection with *C. parvum* and *C. tyzzeri*. Significantly, this effect was observed in two models, immunosuppressed mice infected with the human pathogen *C. parvum* and immunocompetent mice infected with the rodent parasite *C. tyzzeri*. In three experiments performed with customized diet, a significant increase in the elimination of *Cryptosporidium* oocysts was observed in mice deprived of dietary fiber.

The benefits to intestinal health of diets rich in plant fibers are well known [12, 45]. Research on the interaction between the microbiota and the intestinal epithelium has revealed the importance of bacterial metabolites, such as SCFAs originating from the breakdown of plant polysaccharides [6]. Elucidating to what extent this interaction can impact the proliferation of an enteric pathogen could lead to the development of simple “nutraceuticals” capable of mitigating the infection. Dietary supplements would have significant advantages over drugs and vaccines, because they are cheap and do not require refrigeration, a significant advantage for distributing to vulnerable populations such as infants in developing countries. Diet could play a role for controlling cryptosporidiosis as no effective anti-cryptosporidial drugs nor vaccine is available. Moreover, such treatments are unlikely to generate resistant parasites. Although significant, the effect of dietary treatments tested to date on the course of cryptosporidiosis is modest. Clearly, more effective treatments are desirable. Eradication of the infection, however, is not necessarily the most desirable outcome. An intervention which prevents diarrhea, while enabling the host to develop immunity, may be as effective at preventing the long-term consequences of recurrent infant diarrhea [46] than a complete cure. Conceivably, dietary treatments could one day be used to enhance the effect of a drug, when it becomes available, and as prophylactics.

A similar study with the enteric protozoan *Giardia lamblia* concluded that gerbils fed a low-fiber diet were significantly more likely to become infected than animals fed a high-fiber diet [47]. This observation suggests that diet may act directly on the parasite as *Giardia* multiplies extracellulary in the lumen of the intestine. The observed beneficial effect on the course of giardiasis, suggests that dietary treatments may positively affect multiple enteric pathogens.

To maximize the beneficial effect of dietary interventions on cryptosporidiosis, a better understanding of the mechanisms linking diet and parasite proliferation in the intestinal epithelium is needed. The increased severity of certain enteric infections in individuals who eat low-fiber diets can be explained by different mechanisms. A low-fiber diet may increase the abundance of bacteria that degrade the intestinal mucus layer. According to this model [10], infection of enterocytes by enteric pathogens could be facilitated by a depleted mucus layer, one of the main defense mechanisms against pathogens [48, 49]. The effect of diet on parasite proliferation could also be linked to the production of SCFAs or other bacterial metabolites.[50-54]. Given the metabolic dependence of the parasite on host cell metabolites inferred from the annotation of *Cryptosporidium* genomes [55, 56], it is conceivable that bacterial metabolites could affect the parasite’s intracellular proliferation by limiting the availability of essential molecules in the enterocyte.

We previously showed that administration of a probiotic product can aggravate cryptosporidiosis [5]. The prebiotics used here in experiment 4 and 5 are also found in the previously tested probiotics product, but combined with 14 strains of probiotic bacteria belonging to the genera *Lactobacillus, Bifidobacterium* and *Streptococcus* (Supplementary Table S1). The observed mitigating effect of the prebiotics in the absence of probiotic bacteria indicates that the aggravating effect of the probiotics product is caused by probiotic bacteria. Experiments to test the effect of probiotic bacteria, given individually or in different combinations, on the course of cryptosporidiosis may elucidate the mechanisms of interaction between the gut environment and *Cryptosporidium* parasites. Such experiments should combine the analysis of the microbiota and metabolites to identify mechanisms linking diet with parasite proliferation. It is interesting to note that a reduction in the severity of the infection in response to prebiotics occurred regardless of the type of diet consumed. Although an effect on cryptosporidiosis was observed in experiments 4 and 5, the impact on the microbiome was more accentuated in experiment 5 (Table 2). This is likely explained by the fact that both experiment 4 groups already ingested fibers with the diet.

To study the link between diet, intestinal microbiota and the course of cryptosporidiosis, fecal transplant experiments will be needed. Dietary treatments found here to be effective at reducing the severity of cryptosporidiosis in the mouse should also be tested in another model, like the pig [57], to assess the extent to which diarrhea and increased gut motility impacts the effectiveness of the treatment.

## ACKNOWLEDGMENTS

Financial support from the NIAID (grant R21AI125891) is gratefully acknowledged. We thank Albert Tai and the Tufts Genomic core facility staff for high-throughput sequencing.

## SUPPORTING INFORMATION

**Fig. S1. Principal Coordinate Analysis of combined experiments reveals the effect of treatment on the bacterial microbiota.** Each data point represents a fecal sample from one mouse. Samples collected over the entire duration of the experiments are included. A. all experiments (n=212); B. Experiment 4 (antibiotic pre-treatment) excluded (n=162) to de-compress the plot and visualize clustering of samples from the remaining experiments. Symbols indicate experiment as shown in the key. Color indicates treatment as follows: red, no-fiber diet; green, medium-fiber diet; pink, antibiotics followed by prebiotics; black, antibiotics only. File: FigS1.tif

**Fig. S2. Taxonomic classification of microbiota of fecal samples collected on day 0 PI**. Each bar represents one sample collected from one mouse. The experiment number is indicated uppermost. File: FigS2.tif.

**Table S1. Composition of diet and prebiotics.** File: TableS1.xlsx

**Table S2. Bacterial taxa significantly associated with dietary treatment as determined by LDA.** File: TableS2.xlsx

